# Multiple Duration Priors Within and Across the Senses

**DOI:** 10.1101/467027

**Authors:** Darren Rhodes, Anil K. Seth, Warrick Roseboom

## Abstract

Perception can be understood as an active process in which sensory samples are combined with prior expectations to shape perceptual content. A prominent example of the influence of priors on perception is that manually reproduced temporal durations are biased towards the mean of previously experienced durations. However, little is known about how prior expectations are acquired and maintained in environments in which multiple competing cues may indicate whether a given prior should be applied in that specific context. We tested whether human participants could acquire and maintain multiple priors for duration, dependent on the sensory signal in which the duration was presented. Human participants were presented with visual flashes or auditory tones, high or low pitch tones, or white noise versus pure tone audio. In each case, the presented duration on a given trial was drawn from a distribution that was, on average, shorter for tones than for flashes, or vice versa. Our participants’ timing reports were consistent with having acquired distinct duration priors dependent on the sensory signal in which the duration was presented (e.g. auditory or visual). Moreover, this was true whether signals differed across, or within, sensory modality. We account for our findings within a Bayesian framework in which duration priors are iteratively updated depending on determination of a common or distinct origin between successive events. Overall, these results show that the human brain can acquire and maintain multiple perceptual priors based on differences in stimulus properties both within and across the senses.

## Introduction

Humans rely not only on immediate sensation, but also on previous experiences of the world in order to guide perception, decision-making and behaviour. In many studies, behaviour and perception can be described by near-optimal combination of sensory evidence and prior information according to the laws of statistical decision theory [1–4]. Prior knowledge can be acquired over long time scales reflecting stable characteristics of the environment [5–7], and can also dynamically change over shorter time scales as environmental features change [8–12].

Much research in time and timing perception suggests that temporal dimensions of perception follow the same principles [8,9,11–15]. It has long been known that temporal perception is shaped by the context of recent experiences: when presenting a range of inter-mixed intervals, participants’ reports overestimate shorter and underestimate longer durations; this is ‘Vierordt’s Law’ [16,17]. This phenomenon has classically been interpreted as a central tendency effect [18], where participants’ judgments about time are biased towards the mean of the range of presented intervals. More recently, central tendency effects have been interpreted within a Bayesian framework [4,7,8,11,18–22] with the context of recent experience (prior distribution) combined with current sensory evidence (likelihood function) to generate estimations of duration (posterior). Previous investigations of central tendency in duration perception have found evidence for a single temporal prior: participants acquire and maintain information about durations that is shaped by all durations presented, regardless of context or stimulus characteristics [8,15,23].

An important open question, relating these initial findings to more naturalistic settings, is how such processes operate in more complex situations containing several possible sources of information that may systematically vary in their temporal properties. Typical experiments examining the formation of perceptual priors utilise a single context [8,21,23,24], characterised by a constant experimental setup, a single type of stimulus (e.g. visual ready-set-go signals), and/or a single type of response (e.g. manual reproduction). However, well-adapted, naturalistic behaviour requires appropriate attribution of properties, including temporal properties, to their correct sources among many possibilities [25–28].

A recent study [15] investigated how perceptual priors for duration operate when there are multiple potential mappings between expected duration and sensory inputs. Participants were presented with visual stimuli of a range of durations and asked to manually reproduce the presented duration. The stimuli could be presented either to the left or right of a central fixation. The duration of the stimulus presented in a given location (left or right) was drawn from a location-specific distribution of durations such that, for example, left presentations were, on average, shorter and right presentations longer in duration. When presented with this pattern of location-dependent durations, participants’ reports were found to regress to the mean of all presented durations - that is, it didn’t matter that the different locations had different distributions of presented durations, participants’ knowledge of the distribution of temporal properties, their duration *prior*, generalised across location. However, when presenting the same stimulus configuration, but with different *response* modes required for each location (e.g. manual reproduction of right stimulus presentations and vocal reproduction of left) it was found that participants’ reports regressed to the mean of the location-dependent distributions, providing evidence that they had acquired multiple, stimulus(location)-dependent duration priors. These results contrasted with initial findings from our own laboratory [29], in which we found evidence for stimulus-dependent priors, even with a common response mode. Therefore, we here attempt to resolve the apparent empirical inconsistency regarding the conditions under which evidence for the influence of a generalised or stimulus-dependent duration priors can be found.

We used a similar experimental scenario to that described above [15], though with a different task that has previously been found to be highly effective in revealing the influence of perceptual priors in duration perception [9,10]. In this task, on each trial participants are presented with a sequence of four stimuli of a specific sensory signal (e.g., all stimuli in a given sequence could be audio). The intervals between successive stimuli (*ISI*; inter-stimulus interval - the presented durations) are physically identical for the initial three (3) stimuli (two durations) in the sequence, while the final stimulus appears with a timing that is jittered around this duration (*SOA*; Stimulus Onset Asynchrony; see Fig. 1). Participants are required to report whether the final stimulus in the sequence was ‘early’ or ‘late’ relative to what they expected based on the durations between the first three presented stimuli (two durations). From these responses, we estimated the perceived duration for the final (jittered) ISI in the sequence, without the need for the manual (motor timing) responses used in previous work [8,15,21,23,24,30].

**Figure 1.**
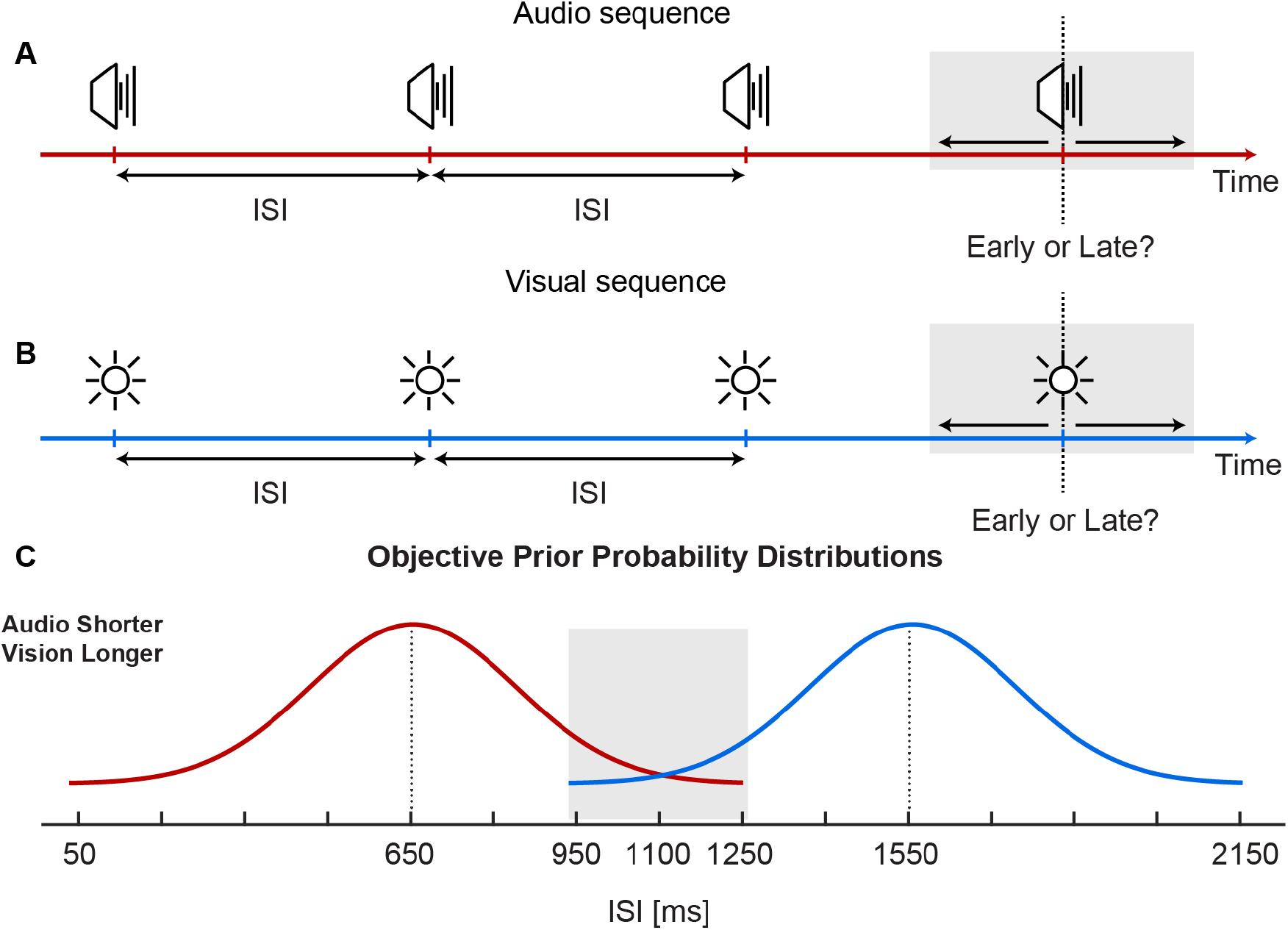
Experiment 1 design and procedure. Participants were presented with sequences of four (A) auditory, or (B) visual stimuli and asked to report whether the final stimulus was earlier or later than expected. (C) In a given block of trials, ISIs were sampled from a Gaussian distribution with a shorter mean for auditory stimuli and longer mean for visual stimuli (and vice versa in a separate session). Long and short mean duration distributions had three overlapping ISIs (950, 1100, and 1250 ms), indicated by the shaded grey area in (C).

In a first experiment, we presented stimulus-dependent distributions of duration (short or long), defined by the ISI, in audio or visual sequences (i.e. in a given block of trials, the distribution of durations for audio sequences was, on average, shorter than that for the visual sequences, or vice versa). In Experiments 2 and 3, duration distributions depended on within-modality differences in presented signal, with short or long duration distributions associated with high or low pitch auditory pure tones (Exp. 2), or with white noise and pure tones (Exp. 3). This method allowed us to seek evidence for the formation of a single, stimulus-generalised duration prior or multiple stimulus-dependent duration priors, indicating the degree to which knowledge-driven biases in human temporal perception generalise or remain dependent on the modality or stimulus type in which durations are defined. A dependency of time perception on sensory modality has often been proposed [31–36], and contrary to previous results [15], we found that priors for duration could be formed dependent on the stimulus properties from which they originate, both across (audio and visual, Exp. 1) and within (two different audio signals, Experiment 3) sensory modality. Critically, the degree to which evidence for stimulus-dependent duration priors could be found was dependent on the featural similarity of the two signals being used. Clear evidence for stimulus-dependent priors was found when the stimuli differed by sensory modality (Exp 1, audio versus visual) or the difference within-modality was highly salient (Exp 3, pure tone versus auditory white-noise) but was not found when the difference was less salient (Exp 2, low versus high pitch pure tone).

We demonstrate that these results are consistent with the output of a Bayesian iterative update model [9,12,37] in which priors are updated and applied following an attribution of common cause between successive samples - an approach similar to that shown to work in the context of causal attribution in other cases of multisensory perception [25,26,28]. This work reconciles the apparent inconsistency between our own and previous [15] investigations into the acquisition of duration priors by providing a framework that demonstrates how apparent source attribution can produce stimulus-dependent or stimulus-general updating of perceptual priors for duration.

## Results

### Experiment 1: Multiple stimulus-dependent priors defined by sensory modality

To determine whether human participants could acquire and maintain two stimulus-dependent priors, in an experimental block we presented participants with sequences of four auditory or visual stimuli on separate trials. On each trial, the first three events were separated by a regular temporal interval, with the last event pseudo-randomly jittered around this interval. Participants reported whether the final event was early or late relative to the expected timing. In a block of 480 trials, intervals between successive auditory tones (ISIs) were, on average, shorter than for visual stimuli, or vice versa (Fig. 1). Trial types were randomly interleaved. To analyse the data, we determined the anisochrony necessary for the perception of subjective on-time responses for each ISI presented, duration distribution (short or long), and modality (audio or visual), by plotting the proportion of “late” responses as a function of the SOA (Point of Subjective Equality; PSE; Fig 2A).

**Figure 2.**
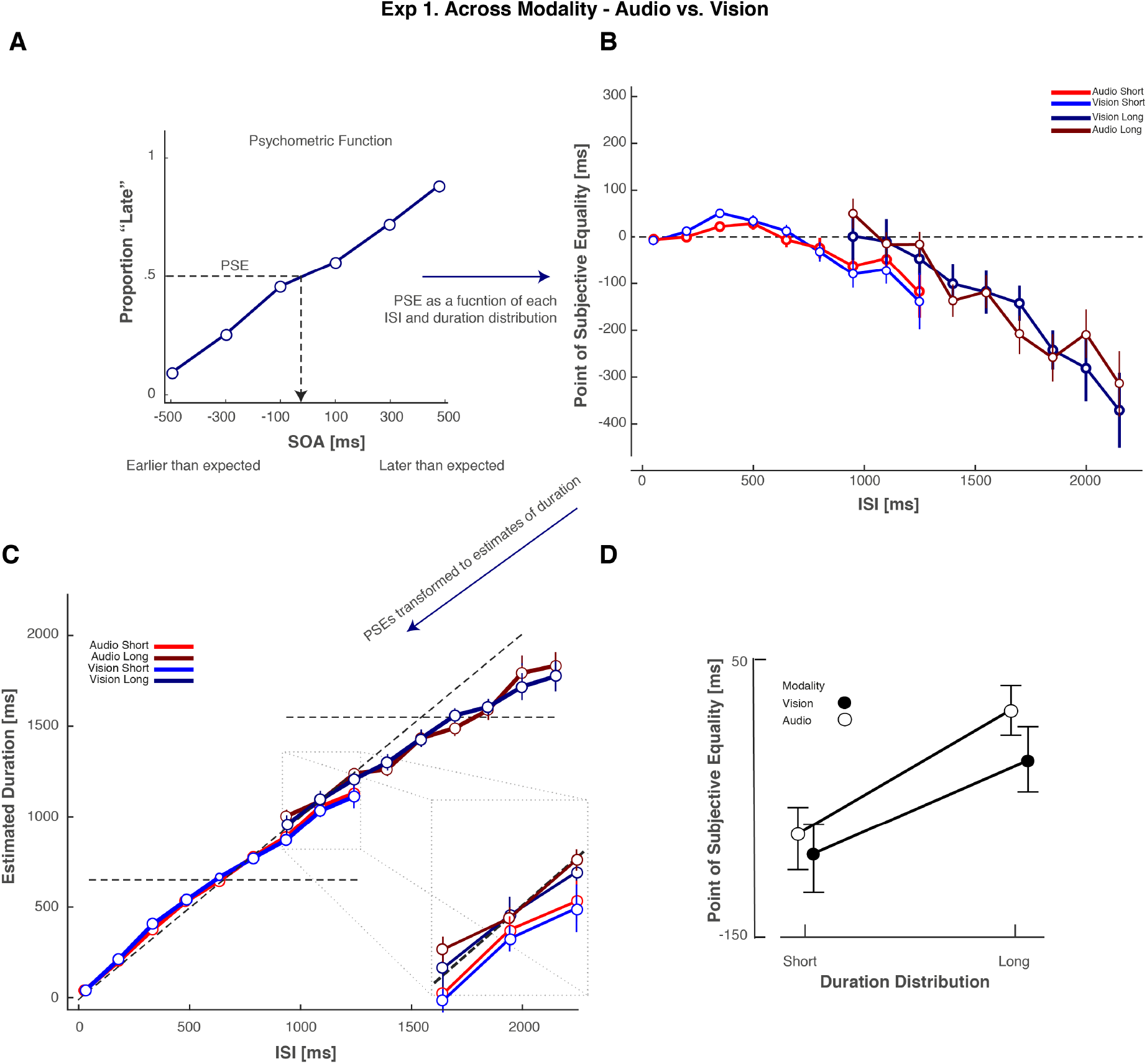
Psychophysical Data Analysis and Experiment 1 Data. (A) Proportion of “Late” responses as a function of SOA for a single ISI. To determine the anisochrony necessary for the subjective report of on-time responses, we performed Spearman-Kärber analyses (See Methods). The PSE is the point at which participants are maximally unsure about whether the stimulus was early or later than expected (dotted line) and therefore provides an estimate of the apparent inter-stimulus-interval. The data presented here is a single example from one subject for the audio short stimulus-dependent duration distribution with an ISI of 1250 ms (B) PSE as a function of ISI for Experiment 1. Each line represents each condition and as such, the different stimulus-dependent duration distributions. An Experimental block consisted of either Audio Short and Vision Long, or in a separate session Vision Short and Audio Long. (C) Estimated durations provided by the Spearman-Kärber Method, with data considered in the (Exp. 1) analyses magnified in each inset. Each line represents a modality-duration distribution contingency. Diagonal dashed lines represent veridical performance on the task, whilst the horizontal dashed lines highlight the mean ISI for short and long duration distributions. (D) Collapsed Point of Subjective Equality (in milliseconds, across ISI) for Spearman-Kärber Method. Error bars represent the standard error of the mean.

As the distributions of ISI for the auditory and visual stimuli contained overlapping values (i.e. duration distributions shared physically identical intervals of 950, 1100, and 1250 ms; see Figure 1C) any difference between them in terms of PSE must be attributable only to the effect of the stimulus (modality) dependency. PSEs were calculated using the Spearman-Kärber Method [38], a method which does not make assumptions about the shape of the distribution that underlies the psychometric function (See Methods for further detail; Fig. 2A). PSE values differed across ISI, such that that they were biased towards the mean of stimulus-dependent ISI distributions, defined by sensory modality.

In this study, both Bayesian and frequentist statistics are reported. Bayes factors assess the strength of evidence for observed data [39–43], and can distinguish between null effects BF_10_ < 1/3, inconclusive data BF_10_ > 1/3 and < 3, and evidence for an effect BF_10_ > 3 [43–45]. Frequentist analyses are also reported for convenience. All statistical analyses were conducted using JASP (Version 9; https://jasp-stats.org). Bayesian analyses used default priors.

As such, a Bayesian repeated measures ANOVA [41,45] with factors ISI (950, 1100, and 1250 ms intervals), duration distribution (short or long) and modality (audio or vision), revealed that a model with the main effects of duration distribution and ISI best described the data BF_10_= 386.156. The model that included modality, duration distribution and ISI described the data with a BF_10_= 119.914, whilst a model including modality and ISI (excluding duration distribution) described the data with a BF_10_ = 0.273.

We additionally submitted the PSE values to a three-way repeated-measures ANOVA with factors of ISI, duration distribution and modality (Fig. 2B). As expected, PSE values (for physically identical ISI) were significantly more positive when presented from the longer duration distribution than when presented from the shorter duration distribution, (Fig. 2BD; Main effect of duration distribution *F*(1, 19)=11.6, *p*=.003, η_p^2^_ = .38). We also found a main effect of ISI *F*(2, 38)=4.6, *p*=.016, η_p^2^_ = .20). The was no evidence to support differences in other main effects or interactions (all *p*>.2). These analyses reveal that the subjective report of stimulus timing was biased towards the mean of a duration distribution dependent on the sensory modality in which it was presented. This provides evidence for prior distributions being built dependent on the modality of presentation: auditory stimuli are biased towards the auditory duration distribution, whilst visual stimuli are biased towards visual duration distribution. As previous investigations of this issue typically rely on estimation of the apparent duration directly, often using manual reproduction, those data depictions usually show duration on the x-axis and reported (reproduced) duration on the y-axis. To facilitate comparison with this previous work [8,11,15,24], in Figure 2C we show the PSE values transformed into perceived duration. This was accomplished by simply adding the obtained PSE to each corresponding ISI in order to provide an estimate of apparent duration [9,10].

### Experiment 2: Can multiple, stimulus-dependent priors be acquired within a sensory modality?

In Experiment 1, we found evidence that our participants could concurrently form multiple distinct duration priors, dependent on the sensory modality in which the stimulus was presented. It has often been suggested that the different sensory modalities may each possess their own, independent temporal processing [33,46]. Therefore, the results of Experiment 1 may demonstrate that this putative independence extends to the acquisition and influence of prior experience of durations on subsequent duration estimation. Alternatively, the influence of multiple distinct duration priors may be possible based on the difference between any sufficiently distinct combination of sensory signals, not just a difference by sensory modality. To test this possibility, in Experiment 2 we used a similar experimental design as in Experiment 1, but this time each duration distribution (short or long mean) was dependent on stimuli that were both auditory, but which were distinct in pitch. Previous work has shown that attributions of commonality or difference within a given sensory modality (e.g. low or high pitch) can be just as effective as differences across sensory modality in multisensory temporal processing [47,48].

As in Experiment 1, we presented participants with sequences of four stimuli, with the fourth stimulus jittered around the expected timing of the final stimulus. On a given trial, the four stimuli could be high or low pitch pure tones, with the ISI between high pitch tones drawn from a long mean duration distribution, and the low pitch tones from a short mean duration distribution, or vice versa. Experiment 2 used a simplified experimental design by comparison with Experiment 1, including fewer levels of ISI, drawn from a uniform rather than Gaussian distribution, and only a single overlapping ISI (Fig. 3; in Exp. 1 there were three shared values). This change in design was done to increase the number of trials collected for each stimulus level-type combination while keeping the overall duration of participation in the experiment session the same. We calculated the PSE for each condition and these are presented as a function of the ISI in Figure 4.

**Figure 3.**
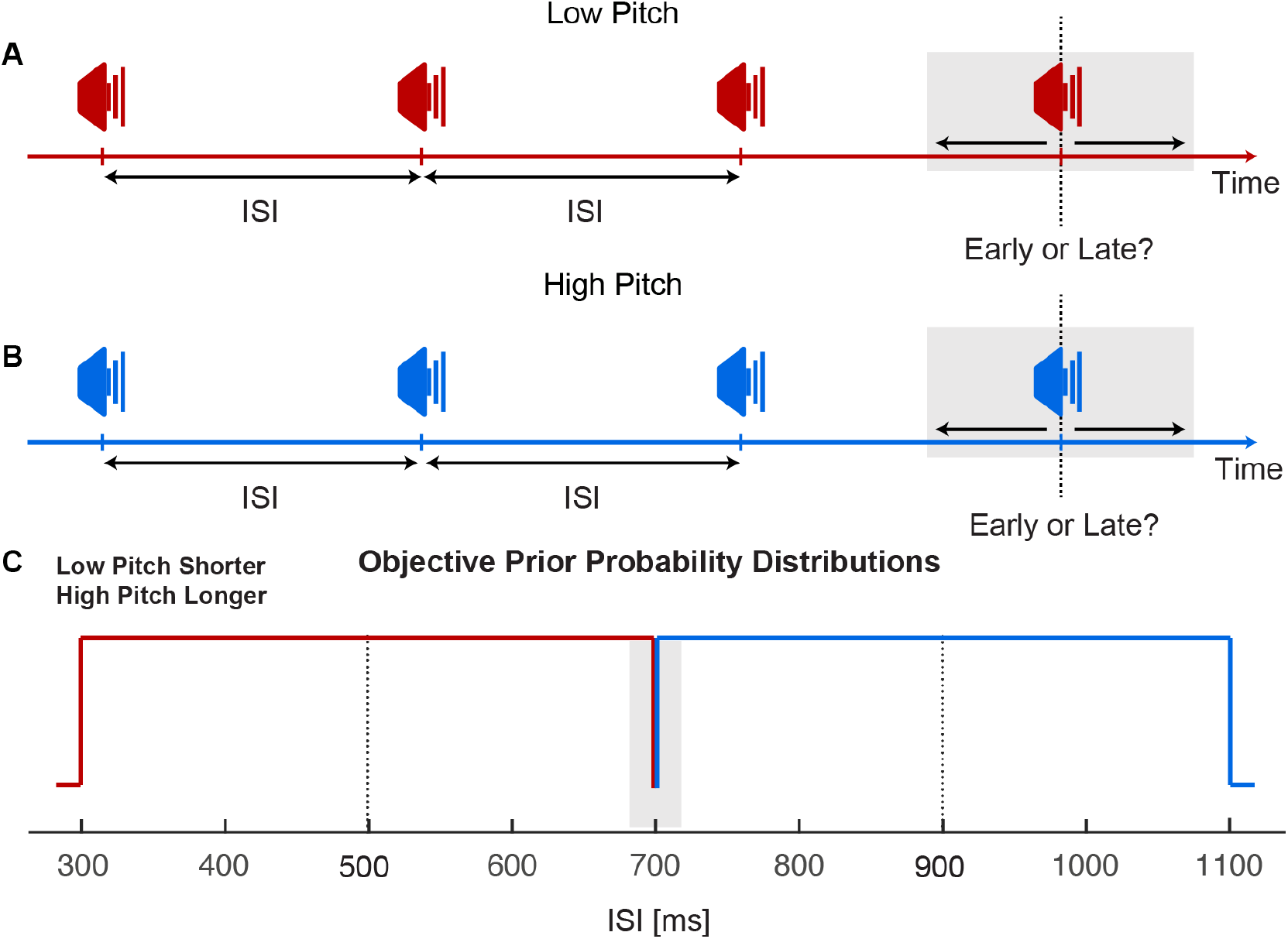
Experiment 2 design and procedure. Participants were presented with sequences of four (A) low pitch, or (B) high pitch tones and asked to report whether the final tone was earlier or later than the expectation. (C) In a given block of trials, ISIs were sampled from a truncated uniform distribution with a shorter mean for low tones and longer mean for high tones (and vice versa in a separate session).

**Figure 4.**
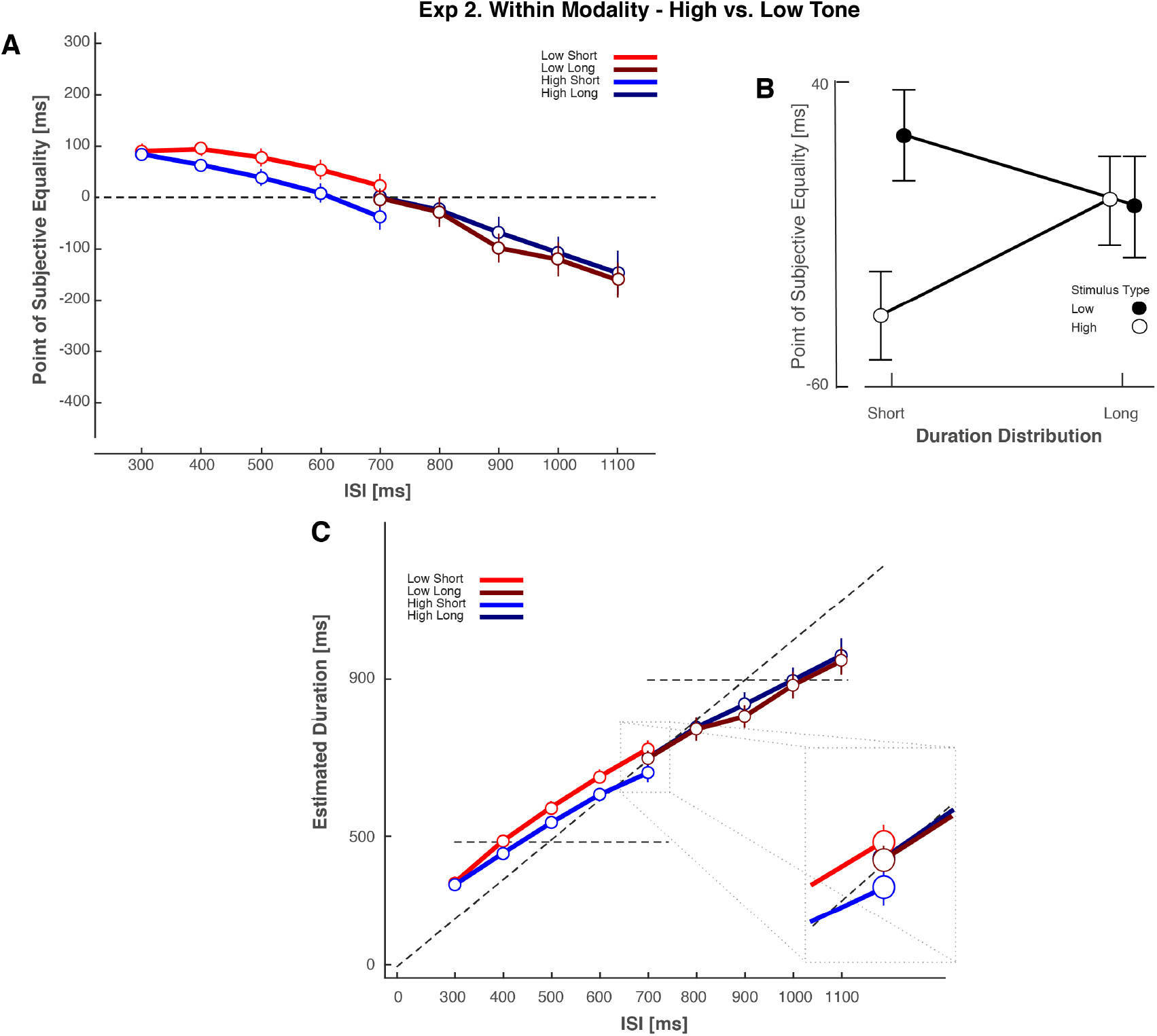
Experiment 2. Data for within-modality priors defined by high versus low tone sequences. (A) PSE as a function of ISI for Experiment 2. Each line represents each condition and as such, the different stimulus-dependent duration distributions. An Experimental block consisted of either Low Short and High Long, or in a separate session High Short and Low Long. (B) Collapsed Point of Subjective Equality (PSE, in milliseconds, across ISI) for Spearman-Kärber Method. Each line represents a modality-duration distribution contingency. (C) Transduced Durations for the Spearman-Kärber Method. Error bars represent the standard error of the mean.

Paralleling Experiment 1, PSEs for each ISI and duration distribution were calculated using the Spearman-Kärber method. The PSE data linearly varied as a function of ISI as (Fig. 4). However, when considering the single overlapping ISI (700 ms) condition, there was no discernible pattern of results consistent with biases towards the mean of stimulus-dependent ISI distributions. Instead, the linear pattern of the data across all ISIs and distributions (short or long) was consistent with a single generalized prior (similar to [15]).

We submitted the PSE values to a two-way Bayesian repeated-measures ANOVA [41,45], with factors of duration distribution (short vs long) and stimulus type (high or low tone; Figure 4AC) for the single overlapping ISI. A Bayesian repeated measures ANOVA revealed that a model with main effects of duration distribution and stimulus type provided evidence consistent with no effect of duration distribution (BF_10_= .251). A model that included just the factor of stimulus type resulted in a slightly larger Bayes Factor (BF_10_= .991) that was insensitive. The model which considered duration distribution alone resulted in a lower Bayes Factor (BF_10_= .246), suggesting the null effects (prior generalization), were driven by the (null) duration distribution factor. In addition, we submitted the PSE values to a 2-way repeated measures ANOVA (Factors duration distribution and stimulus type) and found no evidence that the PSE values from presentations from the longer duration distribution were different from PSE values presented from the shorter duration distribution (Fig. 4AB; Main effect of duration distribution *F*(1, 19)=.312, *p*=.583, η_p^2^_ = .02). The main effect of stimulus type was significant, with low tone stimuli having more positive PSEs *F*(1, 19)=9.024, *p*=.007, η_p^2^_ = .322. The interaction was not significant (*p* >.158). For convenience, we again transduced the PSE values into perceived duration, by adding the ISI to the PSE (Fig. 4C). These results suggest that, unlike in Experiment 1, in Experiment 2 our participants biased their timing reports towards a single, generalized prior over the two distributions of durations defined by high and low pitch auditory stimuli.

### Experiment 3: Evidence for multiple, stimulus-dependent within-modality priors

In contrast to Experiment 1, which provided evidence for multiple stimulus-dependent duration priors when the different duration distributions were presented in distinct sensory modalities, the results of Experiment 2 suggested that duration priors generalise across different stimuli presented from within a single sensory (auditory) modality. This finding may suggest that the acquisition of multiple priors for duration can only occur when the durations are defined within different sensory modalities, consistent with suggestions of separate temporal processing across sensory modalities [33,46]. However, these results may instead indicate that the difference in stimuli in Experiment 2 was simply insufficient to encourage participants to treat the sensory events as distinct and form stimulus-dependent duration priors. Indeed, the results of Roach and colleagues [15] when stimuli differed only by spatial location and not by response mode also failed to produce stimulus-dependent duration priors. One could therefore posit that the degree to which stimuli differ perceptually (in modality, space, time or in some other feature) dictates whether the sensory signals are associated with a common or with disparate sources [25–28,49–51]. To test this premise, we utilized the same experimental paradigm as Experiment 2, but to further encourage our participants to perceive the two different stimulus sequences as being associated with distinct potential sources, we presented sequences of either pure tone or white noise auditory stimuli. We reasoned that pure tones and white noise stimuli, while both auditory, differ more strongly in apparent similarity than high versus low pitch, and so may be more readily treated as unrelated by participants [48,52].

To examine whether stimulus-dependent duration priors could be acquired when stimuli were defined by auditory pure tones and white noise, we analysed the data as in Experiment 2. All experimental methods were the same as Experiment 1 except this time each duration distribution (short or long mean) was dependent on stimuli that were still both auditory, but which were either pure tone or white noise. We calculated the PSEs for each ISI, under each distribution of durations and found a pattern of PSEs similar to the data of Experiment 1 and consistent with stimulus-dependent duration priors within a modality. A Bayesian repeated measures ANOVA [41,45], with factors of duration distribution (short vs long) and stimulus type (pure tone or noise; Fig. 5) revealed that a model with the main effect of duration distribution provided the best evidence BF_10_= 38.764. The model with stimulus type as a factor in conjunction with the duration distribution main effect did not provide as much evidence BF_10_= 9.14, and in tandem with a model including only stimulus type (BF10= .231), suggests that the effect of the prior was equivalent regardless of the stimulus type.

**Figure 5.**
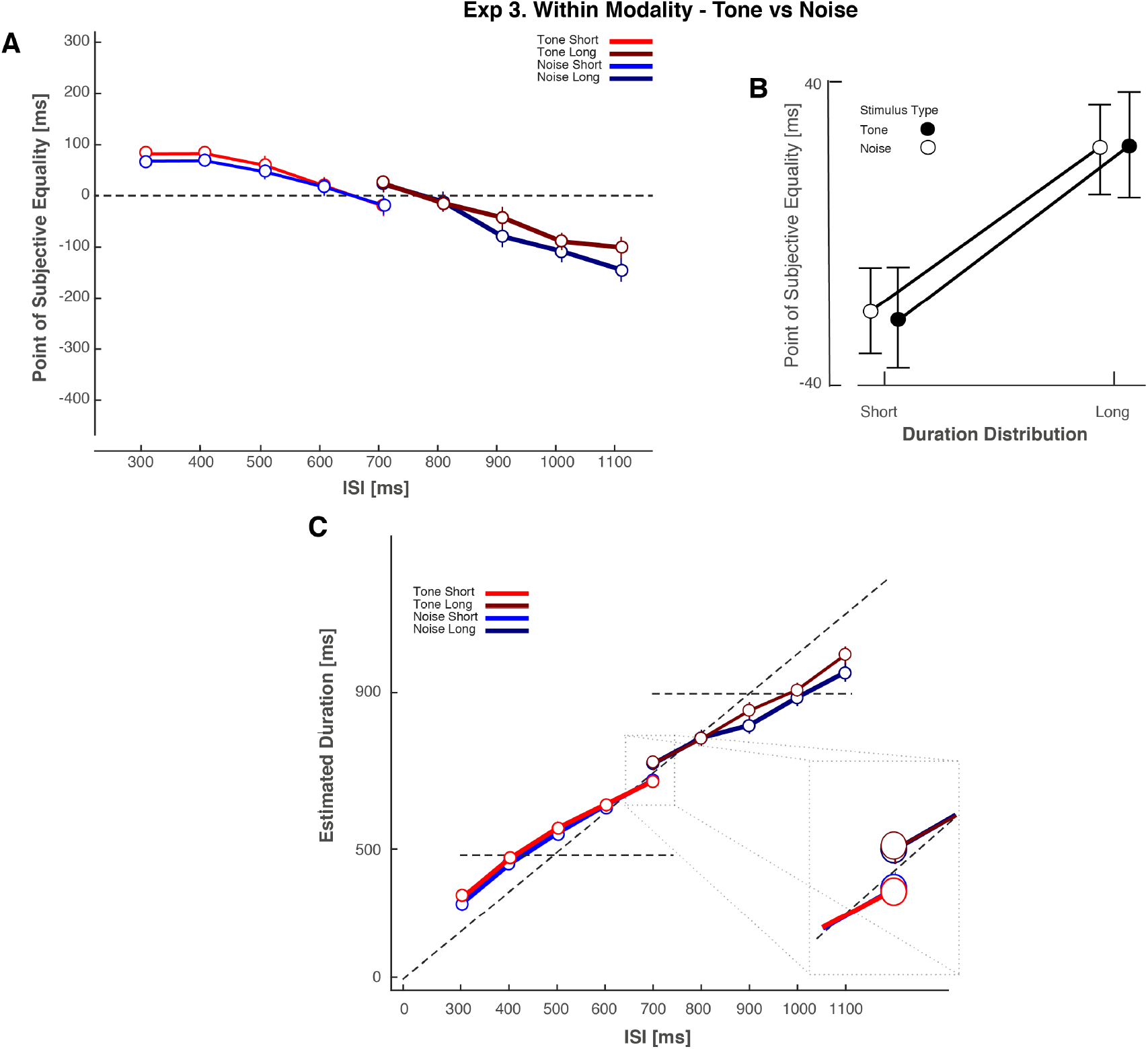
Experiment 3. Data for within modality priors defined by pure tone versus white noise auditory sequences. (A) PSE as a function of ISI for Experiment 3. Each line represents each condition and as such, the different stimulus-dependent duration distributions. An Experimental block consisted of either Noise Short and Tone Long, or in a separate session Tone Short and Noise Long. (B) Collapsed Point of Subjective Equality (in milliseconds, across ISI) for Spearman-Kärber Method Each line represents a modality-duration distribution contingency. (C) Estimated durations provided by the Spearman-Kärber Method. Error bars represent the standard error of the mean.

To provide further evidence that the PSE values were different from each other when presented from different duration distributions, we submitted the PSE values to a 2×2 ANOVA with factors stimulus type and duration distribution. Consistent with Experiment 1 we found PSE values for the shorter or longer duration distributions were statistically different (Fig. 5B; Main effect of contingency *F*(1, 19)=14.7, *p*=.001, η_p^2^_ = .48. The main effect of stimulus type and the interaction were not significant (*p* >.91). These results suggest that participants’ estimates of timing were biased towards the mean of the stimulus-dependent duration distribution, based on the within-modality stimulus characteristics – auditory pure tones versus white noise.

## General Discussion

We investigated whether humans can acquire and maintain multiple priors for duration, depending on the distribution of previously experienced durations associated with a specific stimulus. We found that duration estimates were biased towards the mean of a range of intervals defined by sensory modality (audio or visual; Exp. 1), or other stimulus characteristics when both were presented in the same sensory modality (auditory white noise or pure tone; Exp. 3). By contrast, when the duration distributions were defined by a difference in pitch (auditory low tone or high tone; Exp. 2), we instead found evidence that participants estimates of duration were biased towards the mean of the entire range of intervals – indicating that their prior for duration was generalized across the different presented stimuli. Taken together, these results provide evidence that humans can acquire and maintain multiple concurrent priors for duration when the relevant sensory characteristics are sufficiently different.

Our data provide, to our knowledge, the first direct evidence that exposure to multiple stimulus-dependent distributions of duration lead to changes in *perceptual* decision-making. Previous investigations have revealed that *manually reproduced* durations are biased towards an objective mean of exposed intervals for a single stimulus type and response (manual reproduction) type [8,15,21,23,24]. Here, we reveal evidence for such biases in a purely perceptual task (i.e., not a manual reproduction task). Interestingly, we found that whilst perceptual reports about timing regress towards the mean of a stimulus-dependent duration distribution, the magnitude of the effect is apparently less than in investigations employing manual reproductions [8,15,23,24]. One possible reason for this discrepancy is that perceptual and motor timing judgements are thought to be subserved by different timing systems [53].

Other recent work has also claimed that two priors can be represented for each modality [23]. In that study, participants were presented with two stimuli delimiting an interval, and then asked to manually reproduce the interval. The intervals were sampled from a uniform random distribution, and (in contrast to the design of the present study), were presented in modality-dependent distributions in separate experimental blocks, rather than interspersed within the same block of trials. It is well known that human judgements of auditory and visually presented durations differ in both in their precision (auditory stimuli are more precisely estimated [54]) and bias in estimation (visual stimuli are reported as shorter than auditory stimuli of the same physical duration [55–57]). Given the block-wise design, with each stimulus-type and duration distribution presented in a separate block of trials, along with these known differences in the basic perceptual properties of the used stimuli, this previous study cannot rule out the possibility that differences in the shape of the underlying likelihood functions for auditory and visual stimuli, rather than the influence of two stimulus-dependent duration priors account for the obtained results. By contrast, in our experimental design, we presented both stimulus types and duration distributions interspersed across trials (e.g. auditory-short and visual-long, and the opposite pairings in another experiment block; see Fig. 1). Consequently, our design provides clear evidence for a difference in duration estimation depending on the distribution of durations presented (e.g., Fig. 2), independent of any potential differences in the underlying stimulus properties.

As mentioned in the Introduction, evidence for the concurrent influence of multiple duration priors has been reported recently [15], but only when the duration distributions were dependent on *response format*, rather than stimulus. In contrast to our findings, when the response format was always the same, but duration distributions differed depending on the stimulus (e.g. left location presentations were from a short mean distribution and right from long), these authors found evidence for the influence of a single generalized, rather than multiple stimulus-dependent priors. One may speculate that differing response formats provide clear evidence on which to base stimulus(response)-dependencies and therefore generate stimulus-dependent priors. Perhaps, the mechanism underlying the acquisition and formation of priors generalizes or categorizes stimuli as occurring from a common or separate source, given the strength of available evidence [25–28,50]. In any case, the results presented in this study clearly demonstrate that in addition to response-dependent biases in duration estimation [15], previous experience of stimulus-dependent durations can also bias duration estimation.

### Combining Bayesian causal inference and iterative updating to model stimulus-dependence and generalisation of priors in duration perception

In this paper, we provide further evidence to support previous reports showing that the estimated duration of a given presentation depends on the distribution of durations to which an participant has been previously exposed [8,9,15,21,23,24,58]. Crucially, we additionally demonstrated that this bias in duration estimation towards the mean of the previously observed durations can depend on the stimulus-specific distribution of duration. This stimulus-dependence operates when signals differ both across (audio and vision) and within (auditory white noise and pure tone) sensory modality, but appears to be stronger when the apparent perceptual (featural) difference between stimuli is larger (compare results of Exp, 2 versus Exp, 3). Here, we provide an account able to explain this pattern of results and also reconcile our findings with those previously reported [15], through the application of causal inference models [25,26,28].

Bayesian causal inference has been used to account for behavioural data in many cases in multisensory perception where participants have the possibility of inferring a single common cause or multiple independent causes for sensory stimulation occurring across different sensory modalities (e.g. cross-modal double flash illusion; see [25–28,50,51]). Here, we assume the possibility of (at least) two distinct priors and two separate processes – one which updates priors following each successive sample (trials of the experiment in studies such as presented here), and one which selects which prior to update. By contrast with previous applications of this approach, we use causal inference to determine the apparent causal relation between successive samples (trials) that may be the same or different stimuli (e.g. in Exp 1. Audio and visual), rather than the apparent relationship between concurrently presented multisensory stimuli (such as in, for example, between the auditory and visual stimuli presented in the cross-modal double flash illusion [59,60]).

An example of a model that includes two or more ‘models’ (possible causal structures) is represented by the boxes in Figure 6A. In such systems, an inference process determines the probability of each causal structure. The optimal estimate of any property of interest (stimulus type in our case - audio or visual in Exp 1., low or high pitch in Exp 2, and, auditory white noise or pure tone in Exp 3.) for a given stimulus, *s*̂, is a weighted average of the estimates of two corresponding causal structures. The greater the apparent difference (in space, time, or feature) between two events, the higher the probability that the events originate from two separate sources. Applied to our experiments, consider that the brain is inferring whether successively experienced duration stimuli (successive trials in the experiment) are caused by one general or two distinct sources (audition or vision, auditory low or high pitch, or white noise vs pure tone).

**Figure 6.**
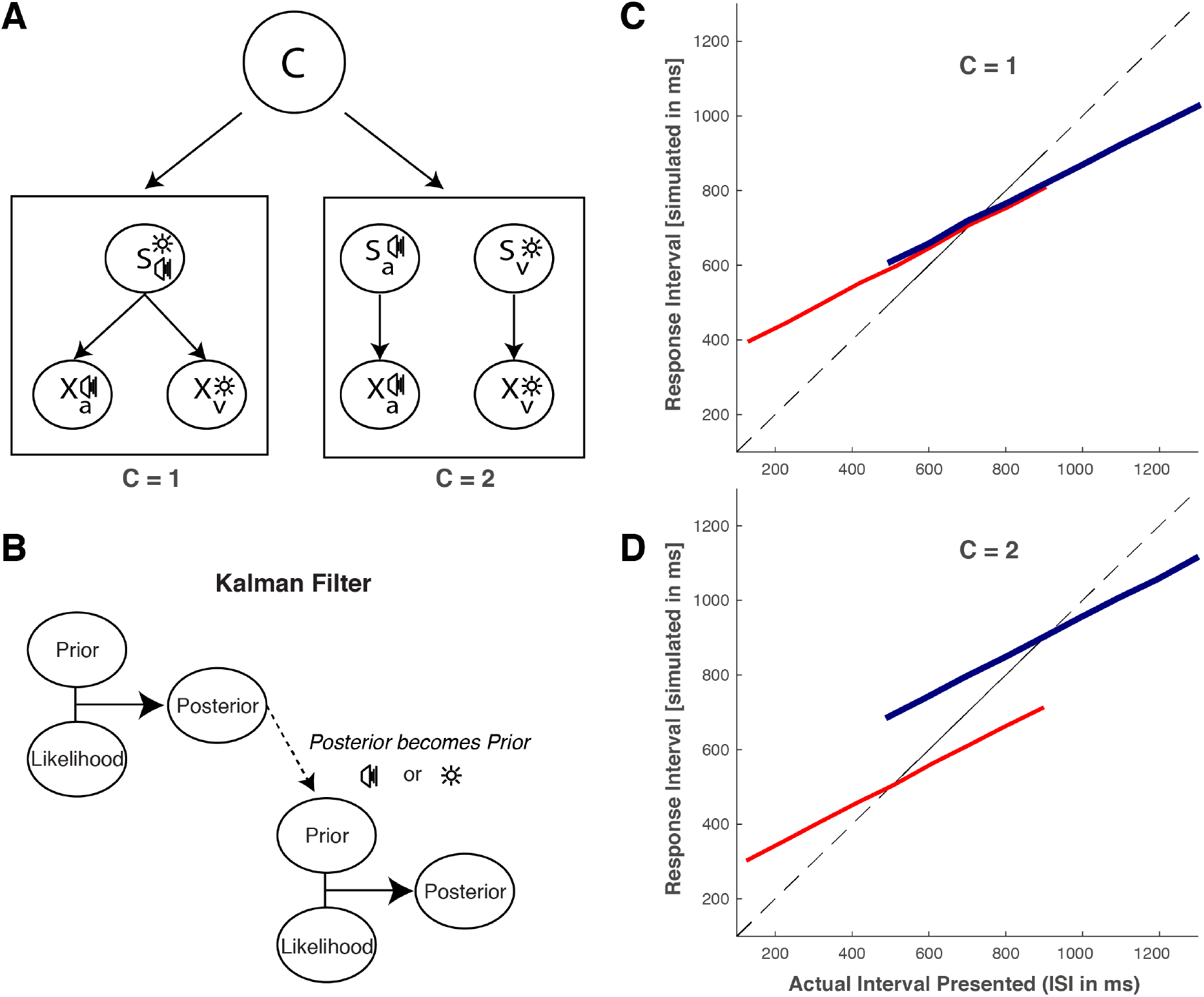
Bayesian causal inference model with a Kalman Filter. (A) Schematic overview of causal inference. Left: One cause may be responsible for both types of sequence. The visual intervals Xv will be the common prior s perturbed by noise σ_v_ and the auditory intervals X_A_ will be the common prior s perturbed by noise σ_A_. Right: Alternatively, differences between distributions and stimulus characteristics might lead to the inference of separate sources of events. The causal inference process infers the probability of a causal structure with a common cause (left; C = 1) and two distinct causes (right; C = 2). The variable C determines which sub-model generates the output. The causal inference layer determines the updating of common or distinct priors using a Kalman filter (B) which iteratively updates prior probabilities about the expected duration of an interval with the likelihood of the current interval. The resultant posterior becomes the prior for the next event. This model qualitatively describes behavioural data from this paper for cases where a single common prior may underlie the data (C; Experiment 2; Fig. 4), or cases where two distinct priors are formed (D; Experiment 1 & 3, Figs. 2 & 5).

We first apply this modelling strategy to Exp. 1, in which sensory signals differed by modality, Figure 6A provides an example of the possible causal structures (see Supplemental Materials for full details of the model). Visual stimuli *X_v_* are perturbed by noise σ_v_, whilst auditory stimuli *X_A_* are perturbed by noise σ_*A*_. The inference process infers the probability of a causal structure with a common cause (left in Figure 6; *C* = 1) and two distinct causes (Right in Figure 6; *C* = 2). The precision and mean estimates of the stimuli interact with the model selection such that situations wherein the different modalities both have low variability will result in model selections that infer two separate causes. By contrast, combinations of stimulus estimates that are more uncertain (i.e. higher noise in σ_v_ and σ_A_) and/or have a smaller difference between means (*X_v_* and *X_A_*) in perceptual space will result in the selection of a single cause. By this method, we can generate cases where successive trials are determined to either relate to the same underlying cause (stimulus; e.g. both visual) or not (e.g. first stimulus visual, second auditory).

The next step is to show how this inference process constrains the formation of priors over repeated experience. Previous ‘summary’ models of duration perception have described prior distributions as inferred from objective sampling distributions [8,21,22,24,61]. A simple sample-by-sample update (trial-by-trial) model is the Kalman filter [9,12,37,62]. A Kalman filter is a recursive filter that estimates the internal state of a linear dynamic system from a series of noisy measurements [62], and can be more accurate than using single measurements [11,12,20,63,64]. Kalman filter-like models have been used widely in the neurosciences, successfully being used to describe central-tendency effects in distance reproduction [12,65], multimodal recalibration [66,67], and sensorimotor control [68]. Applied to our experiments, the algorithm iteratively updates prior probabilities about an expected duration with the likelihood of the current duration estimate, dependent on the strength of available evidence to form stimulus(response)-dependencies (Fig. 6B).

In our model, following determination that the cause of a stimulus is common to that of the previous trial (Fig. 6A; e.g. a visual stimulus presentation, follows a vision stimulus presentation), the iterative-update Kalman filter will operate on the visual prior, combining previous information about visual durations (prior) with the current duration estimate (likelihood). Alternatively, if the outcome of the causal inference process deems the previous trial not to be related to the present trial (e.g. an auditory stimulus following a visual stimulus) then the iterative update process is applied to the other signal, in this case the auditory duration prior. If the causal inference layer cannot effectively infer a difference between one sample and the next, then every trial will be updated under a single, generalized prior. Under this guise, the model can explain both the formation of a single (Fig. 6C) and multiple concurrent priors (Fig. 6D) in temporal perception. Using this approach we may reconcile the discrepancy between our results and that of previous work [15]. When participants can easily determine that stimuli defining the durations (or response modes) are different, these experiments will indicate evidence for stimulus-dependent priors (Fig. 6C; Exp. 1 and 3, here; different response modes in [15]). When the different stimuli (or response modes) are judged to be insufficiently different, evidence will support the influence of a single generalized prior (Fig. 6D; Exp. 2 here, single response mode as in [15]).

An important open question is how the components of Bayesian models such as that described here are represented in the brain. One attractive possibility is that Bayesian inference could be accomplished through probabilistic population coding for perception [69–77] and decision making [78]. This code implies that the natural variability in cortical responses is automatically represented by probability distributions. The computation of likelihoods from such responses is simple and biologically plausible, as just the weighted sum of neural responses [76]. In this approach, prior probabilities are represented by neurons that fire before the presentation of a stimulus. Further, as a prior increases in precision, all neurons with receptive fields for a specific value of a stimulus dimension should increase their gain and fire strongly [74]. Such observations have been reported in the superior colliculus [79] and area LIP [80,81]. Neural oscillations in the brain also offer an interesting way of representing and carrying temporal priors, as the phase of delta-theta band activity could be a plausible neurophysiological mechanism for their implementation [82,83] as well thalamo-cortico-striatal circuits that have also been shown to encode interval timing as well as numerosity [84–87]. However, it still remains unclear precisely where and when the formation of priors, both within and across the senses, occurs in the brain. By addressing such questions, future research will progress towards functioning models of temporal perception that are consistent with human brain anatomy and data [88].

### Summary

Through a series of psychophysical experiments, we show that the human brain can acquire and concurrently maintain multiple perceptual priors, based on differences in stimulus properties both within and across sensory modality. Responses are biased in accordance with simple iterative Bayesian inference, where perceptual estimates are drawn to the mean of previous experience, but depending on a causal inference process to identify which prior to update. Our results extend upon recent findings demonstrating the influence of previous experience on time perception, showing their relevance to scenarios where many different stimuli with different temporal properties must be perceptually processed to guide behavior.

## General Methods

### Participants

Participants were sixty undergraduate and graduate students aged 18-32 from the University of Sussex. Twenty different participants took part in each experiment. All participants reported normal or corrected-to-normal hearing and vision. The ethics committee of the University of Sussex approved the study and all participants provided written informed consent.

### Apparatus and Stimuli

Visual stimuli were generated on MATLAB and displayed on a CRT monitor (resolution of 800 x 600 pixels and a refresh rate of 100Hz) at a distance of 57cm. Audio signals were presented binaurally via Sennheiser HD 280 Pro headphones. Auditory and Visual presentation timing was driven by PsychToolBox [89,90]. Participants responded using the left (“early’) and right (“late”) keys on a standard keyboard.

Auditory signals were the same pitch (frequency of 1Hz) for Experiment 1 and 3, but different for Experiment 2 (3.5Hz vs. .3Hz). Visual events were a luminance modulated Gaussian blob (6 degrees of visual angle; dva). The background was grey. A white fixation circle (.25 dva) was presented centrally with the blob appearing 3 dva above the crosshair. The blob was presented for 2 consecutive frames, approximating a duration of 20ms, whilst auditory signals were also presented for 2 frames with an approximate duration of 20ms.

### Procedure

The task was a binary “early or late” forced choice judgement. Participants sat in a quiet, well-lit room and were presented with a sequence of four auditory or visual stimuli with an inter-stimulus interval (ISI) that was drawn from a distribution with a mean of 650 for short contexts, and 1550 ms for longer ones (Fig. 1). In each context, there were 9 levels of ISI in Experiment 1, and 5 levels in Experiments 2 and 3. The final (4th) stimulus was temporally displaced such that it could appear before or after the expected timing determined by the trial ISI. There were 6 temporal deviations for each of the 9 levels of ISI, which were ±10, 30, or 50% of the trial ISI. In Experiment 1, the temporal deviations were drawn from a uniform random distribution. In Experiment 1 experimental sessions lasted approximately 1 hour each, and participants were presented with sequences where the auditory sequences had a short ISI distribution and the visual sequences had a longer ISI distribution. In the second session, the auditory sequences had a longer ISI distribution, whilst the visual sequences had a shorter ISI distribution. The order of experimental session was counterbalanced across participants with two sessions each.

### Spearman-Kärber Psychophysical Analysis

We analysed the proportion of ‘later’ responses for each anisochrony of the last stimulus, to obtain a distribution for each ISI. In order to determine if a change in the perceived isochrony of stimuli changes due to recent sensory experience, we calculated the *Point of Subjective Equality* (*PSE*) as the anisochrony at which participants are most unsure about whether the final stimulus was presented early or late. Thus, the *PSE* is the time point the last stimulus needs to be presented in order for it to be perceived as being isochronous. The *PSE* is obtained by calculating the first order moment of the difference between successive proportions of responses using the Spearman-Kärber method (see [38,91], for further details of this method). The Spearman-Kärber method is a non-parametric estimate that avoids assumptions about the shape of the psychometric functions underlying participants’ responses. The formulae below are used to estimate the first moment of the psychometric function underlying the data. First, we define the anisochronies of the final stimulus *ANI_i_* with *i*={1, … 15} and *p_i_* with *i*={1, … 15} as the associated proportion of ‘later’ responses. We further define *ANI_0_* and *ANI_16_*= as twice the maximum SOA, whilst also assuming *p0*=0 and *p16*=1, so to be able to compute the intermediate *ANI* between two successive ones

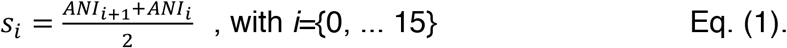

and the associated values of the difference in proportion of responses, taken at and above 0 to monotonize the proportion of responses

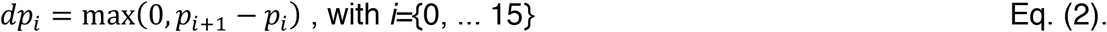

With these indexes, we can express PSE analytically as such:

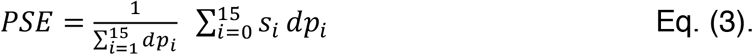

## Ethics Statement

Ethical approval was obtained from the University of Sussex Ethics committee.

## Competing Interests Statement

We have no competing interests.

## Author Contributions

WR, DR, and AKS conceived the study. WR implemented and collected data for Experiment 1. DR implemented and collected data for Experiments 2 and 3. DR analysed the data and performed the modelling. DR and WR wrote the paper and AKS provided critical revisions. All authors gave final approval for publication.

## Acknowledgments

This work was supported by the European Union Future and Emerging Technologies grant (GA:641100) TIMESTORM – Mind and Time: Investigation of the Temporal Traits of Human-Machine Convergence, and by the Dr. Mortimer and Theresa Sackler Foundation, which supports the Sackler Centre for Consciousness Science. AKS is additionally grateful to the Canadian Institute for Advanced Research (CIFAR) Azrieli Programme on Brain, Mind, and Consciousness. The authors also thank Ulrik Beierholm for discussions regarding causal inference.

## Funding Statement

This research was funded by EU FET Proactive grant TIMESTORM: Mind and Time: Investigation of the Temporal Traits of Human-Machine Convergence, with additional support from the Dr. Mortimer and Theresa Sackler Foundation, which supports the work of the Sackler Centre for Consciousness Science.

